# The spatial-temporal dynamics of respiratory syncytial virus infections across the east-west coasts of Australia during 2016-17

**DOI:** 10.1101/2020.11.06.372177

**Authors:** Mark Robertson, John-Sebastian Eden, Avram Levy, Ian Carter, Rachel L Tulloch, Elena J Cutmore, Bethany A Horsburgh, Chisha T Sikazwe, Dominic E Dwyer, David W Smith, Jen Kok

**Affiliations:** NSW Health Pathology - Institute for Clinical Pathology and Medical Research, NSW Health Pathology, Westmead Hospital, Westmead, NSW 2145, Australia; Centre for Virus Research, Westmead Institute for Medical Research, Westmead, NSW 2145, Australia; Marie Bashir Institute for Infectious Diseases and Biosecurity, Sydney Medical School, The University of Sydney, Sydney, NSW 2006, Australia; PathWest Laboratory Medicine WA, Department of Microbiology, Nedlands, WA 6009, Australia; School of Biological Sciences, The University of Western Australia, Crawley, WA 6009, Australia; School of Medicine, The University of Western Australia, Crawley, WA 6009, Australia

**Keywords:** respiratory syncytial virus, molecular epidemiology, Australia, whole-genome sequencing, phylogenetics

## Abstract

Respiratory syncytial virus (RSV) is an important human respiratory pathogen. In temperate regions a distinct seasonality is observed, where peaks of infections typically occur in early winter, often preceding the annual influenza season. Infections are associated with high rates of morbidity and mortality, and in some populations exceeds that of influenza. Two subtypes, RSV-A and RSV-B, have been described, and molecular epidemiological studies have shown that both viruses mostly co-circulate. This trend also appears to be the case for Australia, however previous genomic studies have been limited to cases from one Eastern state - New South Wales. As such, the broader spatial patterns and viral traffic networks across the continent are not known. Here, we conducted a whole genome study of RSV comparing strains across eastern and western Australia during the period January 2016 to June 2017. In total, 96 new RSV genomes were sequenced, compiled with previously generated data, and examined using a phylodynamic approach. This analysis revealed that both RSV-A and RSV-B strains were circulating, and each subtype was dominated by a single genotype, RSV-A/ON1-like and RSV-B/BA10-like viruses. Some geographical clustering was evident in strains from both states with multiple distinct sub-lineages observed and relatively low mixing across jurisdictions suggesting that endemic transmission was likely seeded from imported, unsampled locations. Overall, the RSV phylogenies reflected a complex pattern of interactions across multiple epidemiological scales from fluid virus traffic across global and regional networks to fine-scale local transmission events.

## Introduction

Respiratory syncytial virus (RSV) is a major cause of acute respiratory tract infections in patients of all ages, producing significant morbidity and mortality (Shi et al., 2017). The greatest burden of disease is in children under 1 year old, where it is the most common cause of acute respiratory tract infection, and in this age group is second only to malaria as a cause of death globally (Griffiths et al., 2017; Hall et al., 2009). Importantly, RSV infection in young children may also lead to long-term sequelae such as asthma, chronic bronchitis and obstructive pulmonary disease (Beigelman & Bacharier, 2016; Griffiths et al., 2017). Other populations particularly impacted by RSV include those over 65, and immunosuppressed patients, such as solid organ and bone marrow transplant recipients (Beigelman & Bacharier, 2016; Griffiths et al., 2017).

Despite being discovered in the 1950s, the burden of RSV disease has only recently been appreciated, which is in part due to more reliable methods of diagnosis and detection. The laboratory diagnosis of RSV was initially reliant on viral isolation and the visualisation of characteristic syncytial cytopathic effects, for which its name is derived (Henrickson & Hall, 2007). These techniques were slow and required technical expertise. The most commonly used modalities now include rapid antigen (such as lateral flow immunochromatography and fluorescent immunoassays) and nucleic acid amplification tests (NATs). Molecular assays, including both commercial and in-house NATs, offer increased sensitivity and specificity, as well as the ability to be multiplexed to detect other respiratory pathogens such as influenza, parainfluenza, rhinovirus and human metapneumovirus (Mahony et al., 2007). Rapid test assays can also offer the additional benefit of early diagnosis, which allows for appropriate infection control interventions, rationalisation of unnecessary antibiotic therapy and shorter hospitalisation periods (Henrickson & Hall, 2007). More generally, improved diagnostics and reporting have begun to shed light on the true incidence, seasonal patterns and peaks of activity of RSV. In temperate regions such as the southern major metropolitan areas of Australia, the seasonal peak typically occurs in the late autumn to early winter period (late-March to mid-August) in the months leading into the influenza season (Di Giallonardo et al., 2018; Henrickson & Hall, 2007; Yeoh et al., 2020).

RSV can be divided into two antigenically and genetically distinct subtypes, RSV-A and RSV-B. These may be further divided into genetic groups based on the viral glycoprotein (G gene), termed genotypes, with at least 11 and 23 for RSV-A and RSV-B, respectively. However, recent work has proposed shifting to a genotype classification based on whole genome sequencing (WGS) in order to increase phylogenetic resolution (Ramaekers et al., 2020). Serological studies have shown that the majority of people are infected by age two, and whilst primary infections are typically more severe they are not protective against repeated infection (Griffiths et al., 2017). The spectrum of disease severity associated with RSV infection remains an important yet controversial topic. Studies have made associations between increased severity and particular subtypes/genotypes (Vandini et al., 2017); however these are complicated by both host (Tal et al., 2004) and viral factors (DeVincenzo et al., 2005), as well as their interactions. Molecular epidemiological studies have shown that both RSV-A and RSV-B co-circulate during a season, and often at similar levels (Cattoir et al., 2019; James R. Otieno et al., 2017; J. R. Otieno et al., 2018; Pangesti et al., 2018; Park et al., 2017). Furthermore, for each of these subtypes, a single genotype will tend to predominate, such as with the recent RSV-A ON1-like and RSV-B BA10-like viruses (Di Giallonardo et al., 2018; Eshaghi et al., 2012; Pretorius et al., 2013). Our understanding of the basic molecular epidemiology of RSV has been strengthened by WGS, which, similar to other pathogens, is becoming increasingly common in its application in infectious disease surveillance (Agoti et al., 2014; Dapat et al., 2010; Di Giallonardo et al., 2018). The added resolution from WGS has been particularly useful for elucidating transmission networks at both local (Agoti, Otieno, Munywoki, et al., 2015) and epidemiological scales (Di Giallonardo et al., 2018), as well as for identifying and classifying the introduction and spread of new genotypes (Agoti et al., 2014; Ramaekers et al., 2020).

We recently performed the first genome-scale study of RSV molecular epidemiology in Australia (Di Giallonardo et al., 2018), demonstrating a wide diversity of co-circulating RSV lineages, with limited evidence of strong age and geographical clustering. However, this study was limited to cases obtained from eastern Australia in New South Wales (NSW) through the Western Sydney Local Health District. Here, we expand on these initial investigations to perform a transcontinental study of RSV genomic epidemiology in Australia. We compare RSV strains obtained from Western Australia (WA) to those from NSW on the east coast over an equivalent time period to examine the phylogenetic distribution of strains, and to describe viral traffic between these regions.

## Materials and Methods

### Sample collection and processing

WGS was conducted on residual RSV-positive specimens collected and tested through routine testing at diagnostic laboratories in two major Australian diagnostic laboratories. Specimens were de-identified with basic demographic information collected including age, sex and postcode, as per protocols approved by local ethics and governance committees (LNR/17/WMEAD/128). One hundred and three RSV-A and RSV-B positive cases from WA with a RT-PCR cycle threshold <30 in original screen were collected between January 2016 and May 2017. Sample aliquots (typically, nasopharyngeal swabs in viral transport medium) were transferred to the Institute of Clinical Pathology and Medical Research (ICPMR), NSW, and subsequently extracted using the Qiagen EZ1 Advance XL extractor with the EZ1 Virus Mini Kit v2.0 (Qiagen, Germany). For the eastern Australian RSV cases, 45 previously untyped RSV-positive viral extracts from cases collected during early 2017 at the ICPMR were obtained from storage at −80°C archive. Additional cases from 2016 in NSW were sequenced in a previous study (Di Giallonardo et al., 2018).

### Whole genome sequencing

A previously published approach was used to amplify both RSV-A & RSV-B genomes (Di Giallonardo et al., 2018). In short, RT-PCR was used to amplify four overlapping amplicons (each ~4kb) that together span the RSV genome. The size and yield of each RT-PCR was determined by agarose gel electrophoresis, and the four targets pooled equally. The pooled RSV amplicons were then purified with Agencourt AMpure XP beads (Beckman Coulter, USA) and quantified using the Quant-iT PicoGreen dsDNA Assay (Invitrogen, USA). The purified DNA was then diluted to 0.25 ng/μL and prepared for sequencing with the Nextera XT DNA library prep kit (Illumina, USA). Libraries were sequenced on an Illumina MiSeq using a 300 cycle v2 kit (150 nt paired end reads). Raw paired sequence reads were trimmed using Trim Galore (https://github.com/FelixKrueger/TrimGalore) and then *de novo* assembled using Trinity (Grabherr et al., 2011). RSV contigs were identified by a local Blastn (Altschul et al., 1990) using a database of RSV reference genomes from NCBI RefSeq. The trimmed reads were remapped to draft genome contigs using BBMap (https://sourceforge.net/projects/bbmap/) to check the assembly, and the final majority consensus was extracted for each sample.

### Phylodynamic analysis

The aim of this study was to compare the spatial and temporal dynamics of RSV infection in Australia. To do this, we used a phylogenetic approach to compare the distribution of strains circulating in two major, geographically distinct regions – NSW and WA – representing eastern and western Australia, respectively, during the period January 2016 to June 2017. All WA data, as well as NSW data from the year 2017, was generated within this study. To ensure even sampling across sites, these data were combined with NSW data from 2016 that was obtained from a previous study (Di Giallonardo et al., 2018). To provide additional context, RSV genome data was also sourced from NCBI GenBank where location (country) and collection date (year) was known. The Australian and global RSV genomes were first aligned with MAFFT (Katoh & Standley, 2013), using the FFT-NS-i algorithm followed by manual inspection of gapped regions particularly in the G gene and non-coding regions. The alignments were then trimmed to include only the coding regions, and screened for potential recombinants using RDP4 (Martin et al., 2015) with default parameters that were then removed before analysis. The RSV-A alignment included 1,190 sequences with a length of 15,747 nt, whilst the RSV-B alignment included 1,121 sequences with a length of 15,646 nt. To increase sampling resolution, we also generated comparable datasets using the G gene region only that also included available partial genome sequences from NCBI GenBank that covered the G gene region by at least 300 bp. The final G gene alignments contained 6,603 and 4,300 sequences for RSV-A and RSV-B, respectively, and were both trimmed to the G gene coding region (approximately 960 bp long). The best-fit DNA substitution model for these data was determined with jModelTest (Darriba et al., 2012). Maximum likelihood (ML) trees were then estimated for both RSV-A and RSV-B alignments using RAxML (Stamatakis, 2014) employing the best-fit model, which in both cases was found to be the General-Time Reversible model with a gamma distribution of rates (GTR+G). Support for individual nodes was determined by 1000 bootstrap replications.

### Phylogeographic clustering

From our overall ML trees, it was found that all sequences from 2016 and 2017 were either RSV-A ON1-like or RSV-B BA10-like viruses, therefore we limited further analyses to these specific clusters. To evaluate the geographic structure in the RSV-A and RSV-B trees we used the Bayesian Tip-association Significance (BaTS) program (Parker et al., 2008) with 1000 replicates to analyse the genome datasets. Using BaTS we determined the parsimony score (PS), association index (AI), and maximum clade size (MC) statistics for the location associated with each sequence, specifically focusing on the comparison between eastern and western Australia. To account for other jurisdictions (countries) present in the data, such sequences were assigned to their continent of sampling. This analysis required a posterior distribution of trees, which were obtained using the Bayesian Markov chain Monte Carlo (MCMC) method implemented in BEAST v1.10.4 (Drummond et al., 2012). Here, we used the best-fit DNA model (GTR+G) with a strict clock and a constant population size, as shown to be appropriate previously (Di Giallonardo et al., 2018). All analyses were run for 50 million steps with sampling every 500 steps with 10-20% burn-in. To ensure convergence, three independent runs were conducted and merged to obtain the final set of trees used in the clustering analysis. The maximum clade credibility tree from the Bayesian analyses using BEAST were generated with heights scaled to mean values.

### Local transmission events

In order to identify potential local transmission events (for instance, an outbreak at an institution), we examined both ML and time-scaled Bayesian trees using the genome datasets, for monophyletic groups containing near identical sequences (~99.9% nucleotide identity, less than 10 base pairs different across the genome) sampled up to two weeks apart. Patient demographics and sampling location were then mapped to determine traits associated with these fine-scale phylogenetic groups. The G gene ML trees were then used to validate clustering and to consider additional sources (sampling locations) for local RSV cases.

### Data availability

All sequences generated in this study have been submitted to NCBI GenBank (MW160744 - MW160839). Furthermore, data and material relevant to this study are available from https://github.com/jsede/RSV_NSW_WA.

## Results and Discussion

### Whole genome sequencing of RSV strains from NSW and WA in 2016 and 2017

In order to compare the spatial distribution of RSV strains across eastern (NSW) and western (WA) Australia during the period January 2016 to June 2017, we performed WGS on stored samples collected from community and hospitalised persons presenting with an influenza-like illness. The demographic details of the cohort have been summarised in Table 1. For the NSW cases, no specific sample selection criteria were used except for sample availability from an archived collection of RSV positive residual diagnostic specimens. Most samples from NSW were obtained from children ≤5 years age (51.6%, n=65/126) with a further 20.6% of samples collected from patients ≥65 years (n=26/126). In contrast, samples from WA were selected to represent all age groups. Despite this, similar to NSW, most available samples were collected from the young (62.1%, n=64/103 from patients ≤ 5 years age) and the elderly (17.5%, n=18/103 from patients ≥65 years), reflecting the distribution of infection at both extremes of age. The distribution between sexes was approximately equal for both study sites (Table 1).

**Table 1.** Demographic details of NSW and WA RSV cohorts.

| Age | NSW | WA | Sex | NSW | WA |
| --- | --- | --- | --- | --- | --- |
|  | Number of samples (%) |  |  | Number of samples (%) |  |
| ≤5 | 65 (51.6%) | 64 (62.1%) | Male | 66 (52.4%) | 48 (46.6%) |
| 5 - 65 | 35 (27.8%) | 21 (20.4%) | Female | 60 (47.6%) | 55 (53.4%) |
| ≥65 | 26 (20.6%) | 18 (17.5%) | Total | 126 (100%) | 103 (100%) |
| Total | 126 (100%) | 103 (100%) |  |  |  |

Virus genomes were successfully extracted and sequenced from 63 of the 103 respiratory samples collected in WA during 2016 and 2017, and 32 of the 45 NSW cases collected in 2017 (total new genomes n=96, where one case from NSW was an RSV-A/B mixed infection). These data were combined with existing genomes from NSW during 2016 (n=93) generated previously (Di Giallonardo et al., 2018), bringing the total number of Australian genomes for analysis across both states in 2016 and 2017 in the current study to 189. The breakdown between sampling location and RSV subtypes determined by WGS shows that both RSV-A and RSV-B subtypes were present in both NSW and WA populations (Figure 1). In NSW, there was a higher proportion of RSV-B strains (63%, n=80/126), which was consistent across the entire study period. Furthemore, in NSW, the peak in RSV activity typically occurs in May-June each year (NSW Health influenza surveillance data), however most of our genome sequences in 2016 were from isolates in August to October that year; that is, they were from the later part of the RSV season. In WA, there was even representation of RSV-A and RSV-B in the WGS data across 2016 with an increase in the relative proportion of RSV-A during 2017. While our sampling for WA was intentionally even with regards to time, subtype, age and setting, the predominance of RSV-A during 2017 was also observed by the initial diagnostic testing where the used resolves RSV subtypes. Specifically, with this lab testing data a transition from RSV-B to RSV-A was observed between the 2016 and 2017 seasons in WA (data not shown). These results are consistent with other molecular epidemiological studies globally that show both RSV-A and RSV-B subtypes co-circulate with shifting predominance across seasons (Di Giallonardo et al., 2018; Luo et al., 2020; Razanajatovo Rahombanjanahary et al., 2020; Yun et al., 2020)

**Figure 1.**
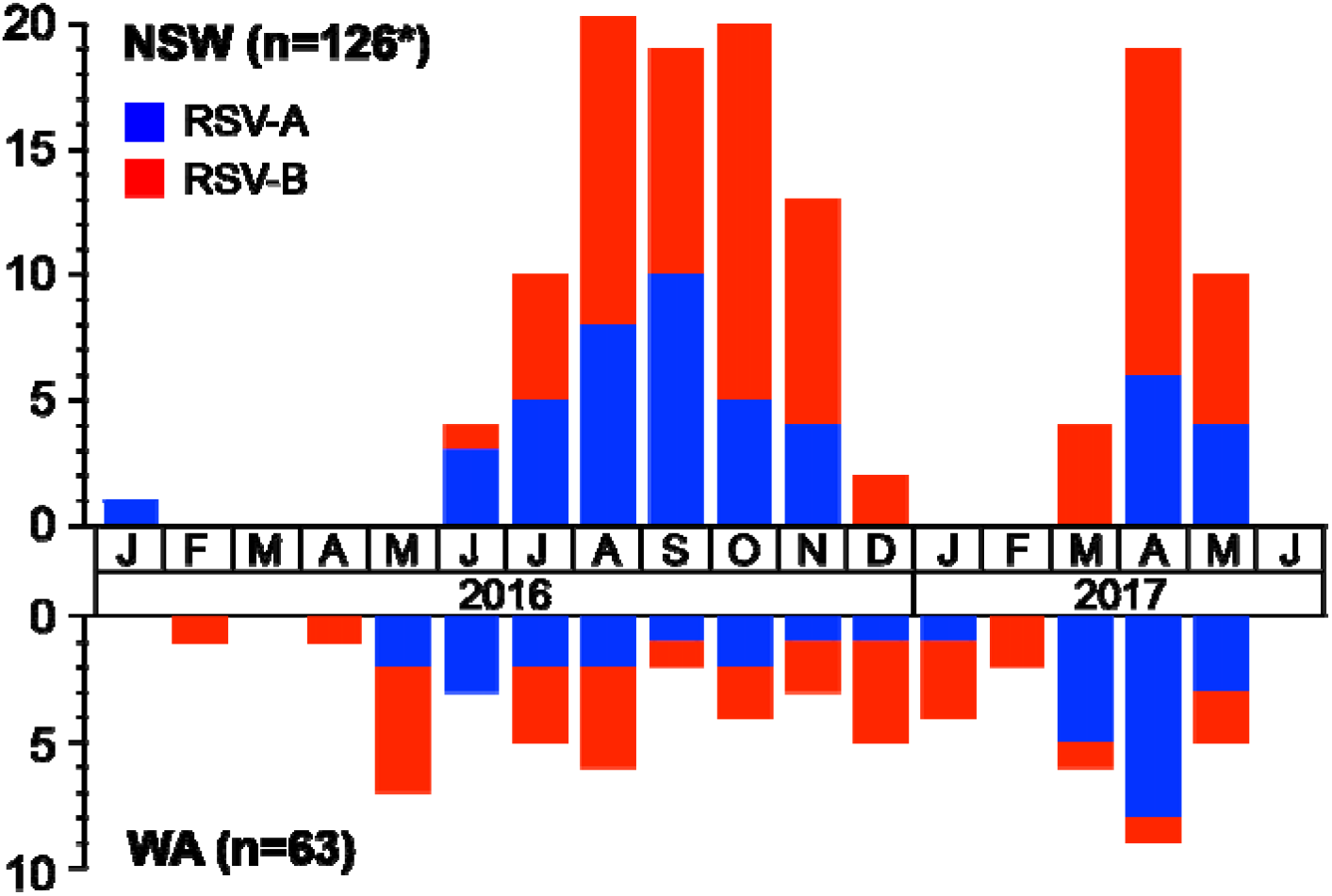
Whole-genome sequencing of respiratory syncytial virus (RSV) in Western Australia and New South Wales between January 2016 and June 2017 by month. The y-axis represents counts of genomes sequenced. The x-axis represents months when the samples were collected, with year shown underneath. RSV-A and RSV-B subtypes have been coloured blue and red respectively, as per key provided. In total, over the period of investigation, 189 RSV genomes were compared including 63 from WA and 126 from NSW. WA samples were selected pre-sequencing to cover both subtypes during seasonal and inter-seasonal periods and therefore do not necessarily reflect the subtype or temporal distribution of RSV in WA. The asterisk indicates that of the 126 NSW genomes, 93 genomes from 2016 were generated in a previous study (Di Giallonardo et al., 2018).

### Phylodynamics of RSV infections in NSW and WA in a global context

The WGS data from NSW and WA were aligned and compared to global genome references to provide genetic context for local strains. Phylogenetic analysis using a maximum likelihood approach was then employed to examine the diversity of RSV-A and RSV-B strains (Figures 2 & 3). This analysis revealed that for each subtype, regardless of location, a single genotype was predominant. For RSV-A, recent viruses from both NSW and WA were derived from the ON1 lineage (Eshaghi et al., 2012), which has been the predominant RSV-A genotype globally since it first emerged in 2011 (Agoti et al., 2014; Di Giallonardo et al., 2018; Eshaghi et al., 2012; Tabatabai et al., 2014)(Figure 2). Similarly, the RSV-B phylogeny showed that circulating viruses were mostly of the BA10 lineage (Figure 3). From the limited available sequence data on GenBank, it appears that the BA10-like viruses are circulating globally, however, there are few published molecular epidemiology studies to support the suggestion they are predominant. It is clear however, that in Australia, in both NSW and WA, that this has been the major RSV-B genotype since at least 2014 (Di Giallonardo et al., 2018).

**Figure 2.**
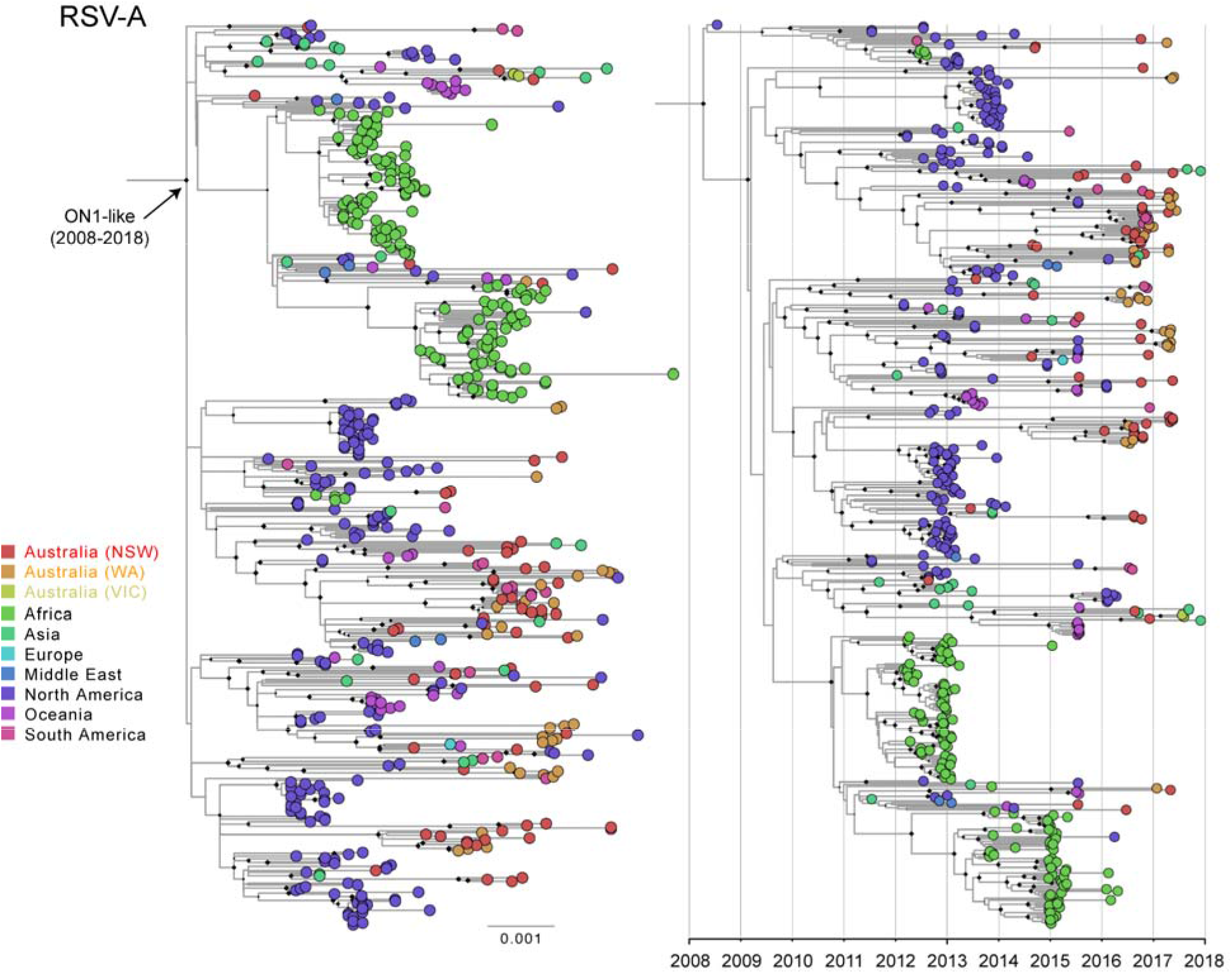
Phylogenetic analysis of respiratory syncytial virus (RSV) A strains circulating in Western Australia and New South Wales between January 2016 and June 2017. The global phylogenies were first estimated using alignments of complete genome sequences of RSV-A (n=1,190) strains circulating since the late 1970’s, however for clarity, only the recently circulating RSV-A ON1 lineage has been shown (n=499), which is defined by the specific branch marked with an arrow. The panel on the left shows the tree estimated using a maximum likelihood (ML) approach, while the tree on the right shows the same phylogeny estimated using the time-scaled Bayesian approach. The taxa, shown as small circles, have been coloured by sampling location as per the key provided. The red and orange coloured circles sequences sampled in NSW and WA, respectively. Diamonds at nodal positions indicate branching support with bootstrap replicate values >70% and posterior probability values >0.9 for the ML and time-scaled Bayesian trees, respectively. The left sided scale bar represents the number of substitutions per site, while the right is scaled to time (years).

**Figure 3.**
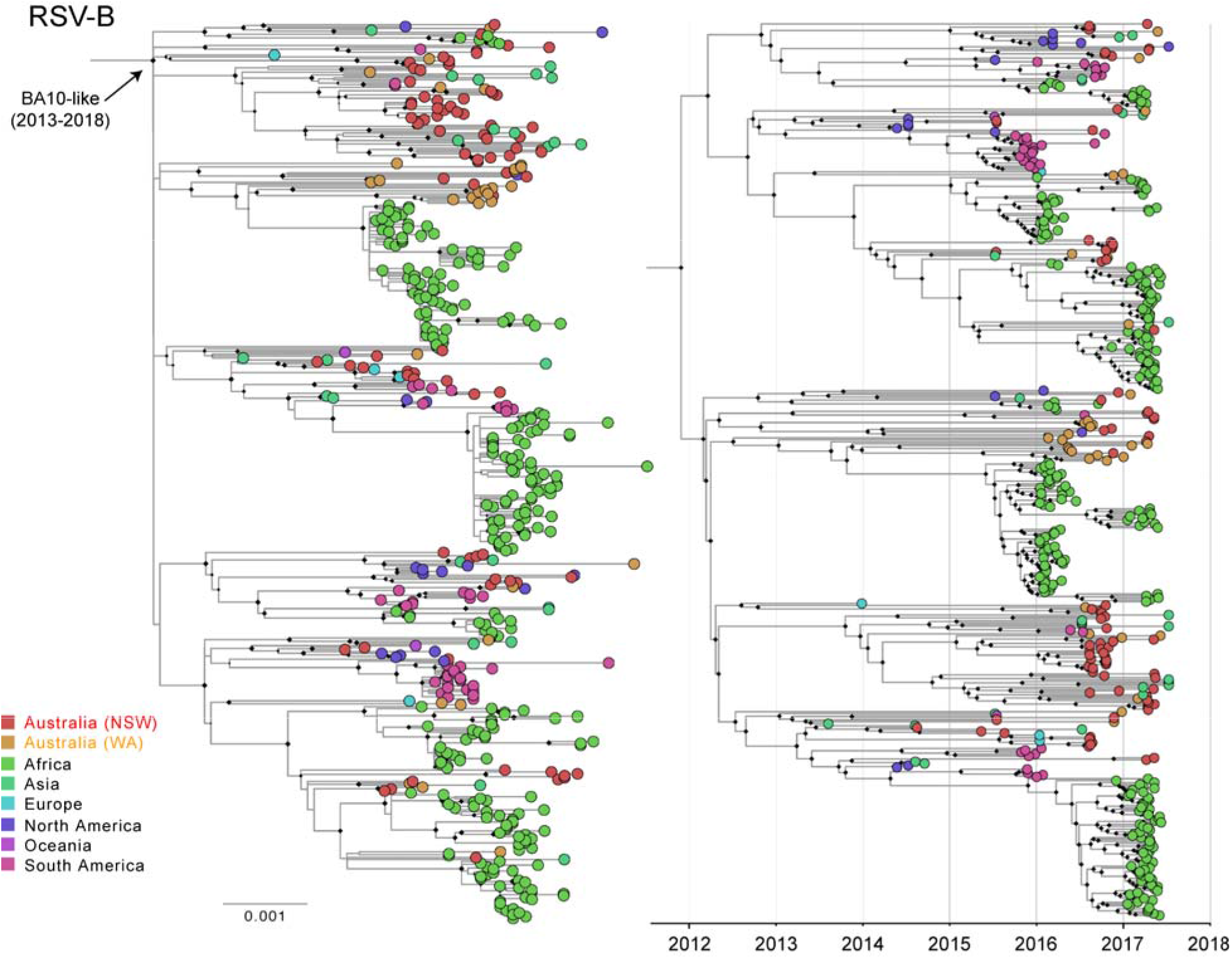
Phylogenetic analysis of respiratory syncytial virus (RSV) B strains circulating in Western Australia and New South Wales between January 2016 and June 2017. The global phylogenies were first estimated using alignments of complete genome sequences of RSV-B (n=1,121) strains circulating since the 1960’s, however for clarity, only the recently circulating RSV-B BA10 lineage has been shown (n=470), which is defined by the specific branch marked with an arrow. The panel on the left shows the tree estimated using a maximum likelihood (ML) approach, while the tree on the right shows the same phylogeny estimated using the time-scaled Bayesian approach. The taxa, shown as small circles, have been coloured by sampling location as per the key provided. The red and orange coloured circles sequences sampled in NSW and WA, respectively. Diamonds at nodal positions indicate branching support with bootstrap replicate values >70% and posterior probability values >0.9 for the ML and time-scaled Bayesian trees, respectively. The left sided scale bar represents the number of substitutions per site, while the right is scaled to time (years).

Across both RSV-A and RSV-B phylogenies there does not appear to be strict spatial clustering when comparing viruses from NSW and WA (Figures 2 & 3). That is, the viruses from WA and NSW do not form monophyletic groups. Such monophyletic clustering would not be expected based on what is known for other respiratory pathogens - such as influenza - which are characterised by high levels of gene flow and viral traffic at global scales (Rambaut et al., 2008; Vijaykrishna et al., 2015). As such, we observed multiple co-circulating sub-lineages distributed across the entire diversity of RSV-A/ON1-like and RSV-B/BA10-like viruses (Figures 2 & 3). However, within these sub-lineages the viruses often formed clusters based on sampling location, therefore, local spatial structure was apparent in our analysis. To explore this formally, we conducted a clustering analysis with BaTS, and a posterior set of trees estimated using a time-scaled phylogenetic analysis in BEAST. We limited the analysis to the RSV-A/ON1 and RSV-B/BA10-like viruses, and the maximum clade credibility tree for each RSV subtype were found to be congruent with the ML trees (Figures 2 & 3). Using the posterior set of trees we then measured the degree of clustering based on sampling location (Table 2).

**Table 2.**
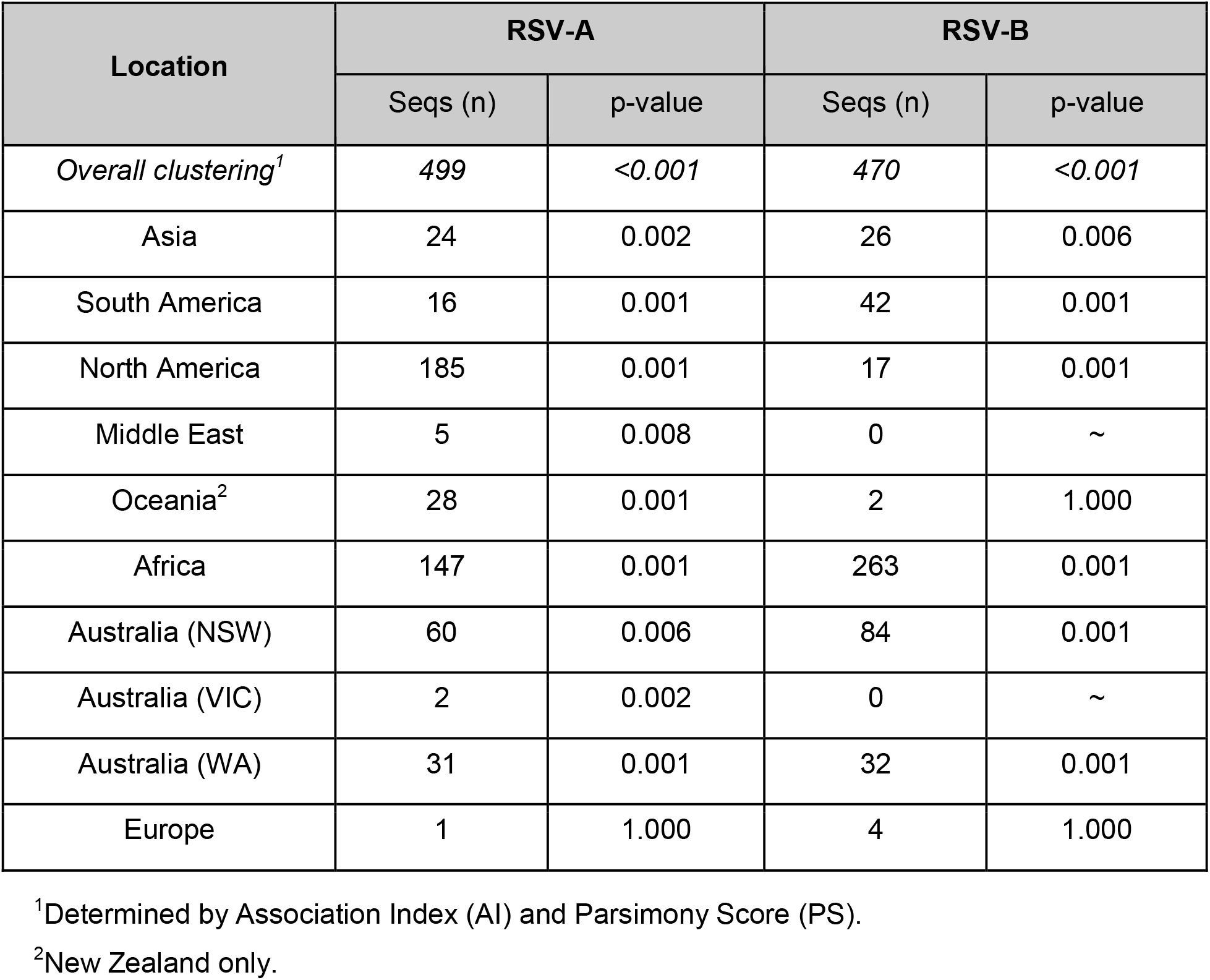
Phylogeny-trait association test for RSV in Australia and other regions globally.

When sequences were grouped by broad geographical regions, specifically, continents and the two Australian states NSW & WA (and VIC for RSV-A data), phylogeny-trait association tests indicated a significant pattern of overall geographical clustering (p-values <0.001) as measured by AI and PS scores (Table 2). For individual locations, most were found to cluster with significant scores. This includes Asia, the Americas, the Middle East, Africa, Oceania, NSW, WA and VIC for RSV-A viruses (all p-values <0.008) and then similarly all for RSV-B where the sampling was sufficient (n >4 sequences). This analysis demonstrates that, at larger epidemiological scales, viral lineages are often imported and established locally. We note however, that this analysis is biased for regions that have been highly sampled, as highlighted by the American and Kenyan viruses, which are mostly from one specific city and/or hospital (Supplementary Figures 1 & 2) (J. R. Otieno et al., 2018). Our sampling of NSW and WA strains is similarly constrained, except that our sampling sites are both major diagnostics hubs which perform testing across wide geographic regions that are representative of large and diverse populations (Di Giallonardo et al., 2018), and in our comparison the period of sampling was equivalent. Despite this and any apparent geographical clustering, the phylogenies both reflect a lack of data from many jurisdictions globally, and our insights into global and local viral traffic will remain hampered until these genome sequences become available. To address this, we also performed ML phylogenetic analysis on a more well-sampled dataset derived from sequences of the G gene region only (Supplementary Figures 3, 4 & 5). While this analysis improved the overall location sampling, particularly for European and Asian sequences, there was less phylogenetic resolution, therefore a trade-off exists between using the less sampled, higher resolution whole genome data versus the more well sample partial genome (G gene) sequences available on NCBI GenBank (Supplementary Figure 3). Regardless, as shown below, this data can shed light on the possible sources and global transmission networks for RSV.

### Lineages and local clusters

A detailed examination of individual lineages (Supplementary Figures 1 & 2) confirmed that most only contained viruses from either state. There were some instances where individual viruses from NSW were nested within diversity solely comprised of WA strains, and vice-versa, and which may represent viral migration events between the jurisdictions. The overall pattern suggests low levels of mixing between NSW and WA, perhaps due to the large geographic area of each state and the distance that separates them. Moreover, we had suggested previously that some evidence of viral persistence across seasons was present in NSW (Di Giallonardo et al., 2018). Here, we found no direct link between the 2016 and 2017 seasons in both jurisdictions and that viral diversity is likely maintained and seeded from imported yet unsampled locations. An examination of the better-sampled G gene phylogenies (Supplementary Figures 4 & 5) did not clarify potential additional sources, except for sporadic examples where NSW or WA were clustered genetically and temporally with Asian or European strains. Importantly, we also found examples where two Australian sequences clustered in the G gene phylogeny but were then sufficiently different at a genome scale to not be classified as clusters such as seen with the NSW and WA strains WM1079A/2016-08-14 (MH760625) and PW3375488K/2016-08-09 (MW160759) (Supplementary Figures 1 & 4). This is consistent with other studies that have shown that local persistence makes a minimal contribution to the diversity of RSV strains in any given area (Agoti, Otieno, Ngama, et al., 2015; Zou et al., 2016). Rather, viral importation and global mixing of strains seem to be the major driver of RSV diversity and their sources. Despite the data suggesting limited east-west viral traffic across the Australian continent, the Sydney-Perth air route remains one of the busiest in the country, nor can we discount other regions within Australia such as the tropical North, where persistence of viral lineages may occur as important sources of viral diversity. Indeed, from our understanding of influenza virus in Australia, the synchronized dissemination of viral strains across the country is driven by both multiple introductions from the global population and strong domestic connectivity (Geoghegan et al., 2018). Increasing the breadth of WGS data for RSV cases nationally would assist in confirming what domestic viral traffic networks exist, particularly between tropical and temperate regions, as well as, between major population centres on the east coast such as Sydney and Melbourne.

Next, we examined genetic clusters defined by high genetic identity (<10 bp difference across the genome) and similar sampling periods (collection dates within two weeks) to consider what features such as patient age and sampling location define them. For RSV-A, we identified 8 clusters (Supplementary Figure 1 & Table 1) and for RSV-B we identified 10 clusters (Supplementary Figure 2 & Table 2). In this study 75% (n=6/8) and 80% (n=8/10) of RSV-A and RSV-B clusters, respectively could be linked to common localities and/or institutions, including individual hospital wards or emergency departments. While two of the RSV-B clusters were found to be repeat samples of the same patient (Supplementary Table 2), overwhelmingly, these clusters of genetically and temporal related sequences most likely represent fine-scale transmission events.

## Conclusions

In summary, our analysis has identified a number of features of RSV epidemiology and patterns of spread including i) the co-circulation of both major subtypes: RSV-A and RSV-B; ii) a single genotype for each subtype predominates each season; iii) multiple distinct sub-lineages of each genotype will co-circulate and which are associated with regional and local clustering/outbreaks, iv) little viral mixing across the east-west coasts of Australia despite apparent overall geographic clustering, v) that genetically and temporal related sequences most likely represent fine-scale transmission events such as institutional outbreaks and vi) that whole genome sequence is required and encouraged over partial G gene sequencing for elucidating clusters and transmission pathways. Taken together, this presents a complex phylodynamic pattern with globally circulating diversity with viral mixing across different regions, yet, finer-scale patterns revealing multiple endemic sub-lineages and clusters consistent with local transmission events and outbreaks. This highlights the connections of genomic data across multiple epidemiological scales and further strengthens the need for much greater sampling of RSV, not just here in Australia, but globally.

## Acknowledgments

We thank all the members of the Virology and Microbiology teams at NSW Health Pathology - ICPMR in Westmead, NSW and the PathWest QE2 Medical Centre laboratories in Perth, WA for all their contributions towards processing the diagnostics specimens used in this study.

