## Supplemental Figure 3 for "The spatial-temporal dynamics of respiratory syncytial virus infections across the east-west coasts of Australia during 2016-17"

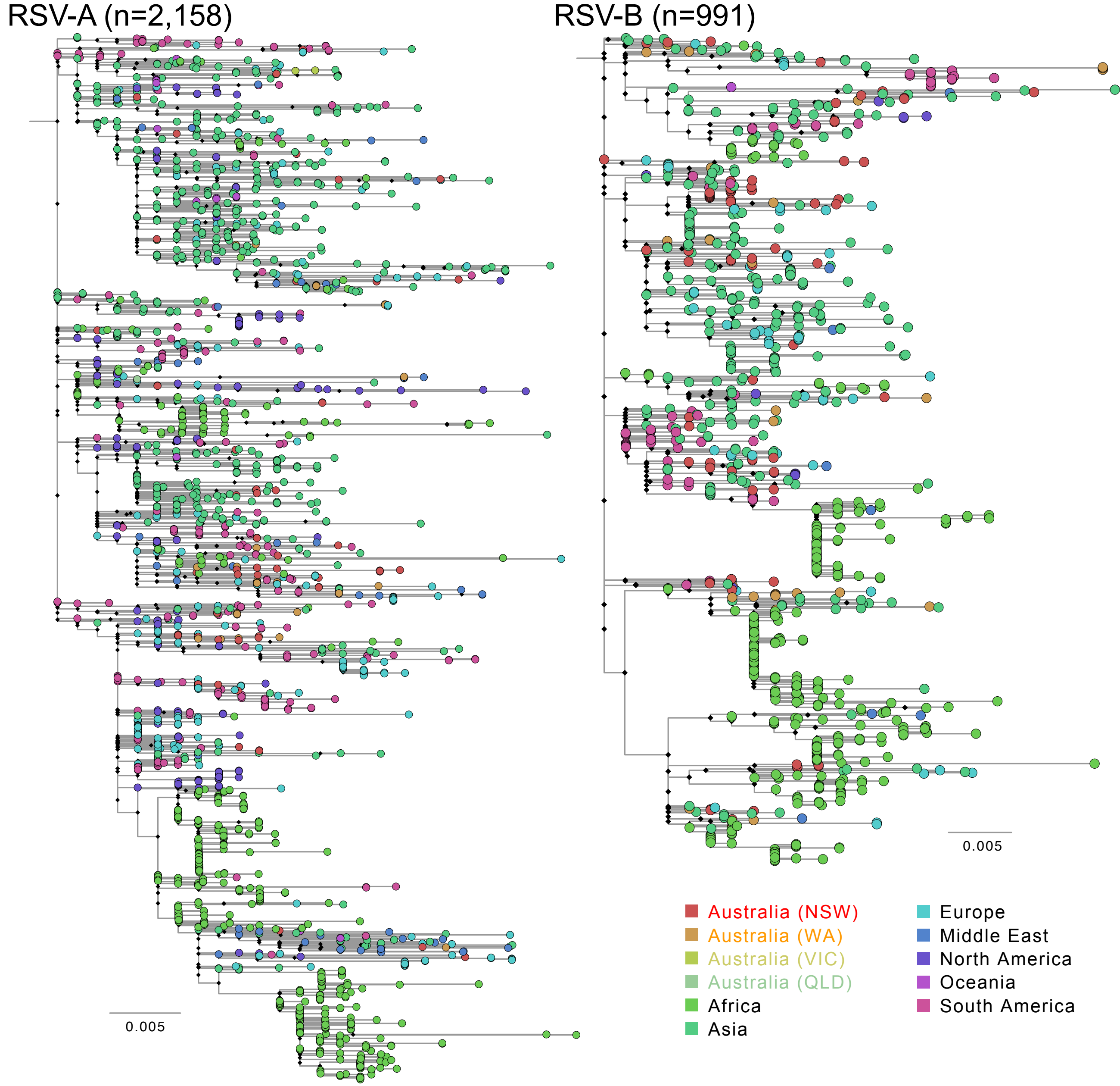


**Supplementary Figure 3. Maximum likelihood phylogeny of respiratory syncytial virus (RSV) strains circulating in Western Australia and New South Wales between January 2016 and June 2017.** The phylogenies first estimated using alignments of complete and partial G gene sequences of RSV-A (n=6,603) and RSV-B (n=4,300) strains circulating since the 1950's, however for clarity, only the recently circulating RSV-A ON1 and RSV-B BA10 lineages are shown (RSV-A and RSV-B on the left and right panels, respectively). The taxa, shown as small circles, have been coloured by sampling location as per the key provided. The red and orange coloured circles sequences sampled in NSW and WA, respectively. Diamonds at nodal positions indicate branching support with bootstrap replicate values >70%. Scale bars represent the number of substitutions per site.
