## Supplemental Figure 4 for "The spatial-temporal dynamics of respiratory syncytial virus infections across the east-west coasts of Australia during 2016-17"

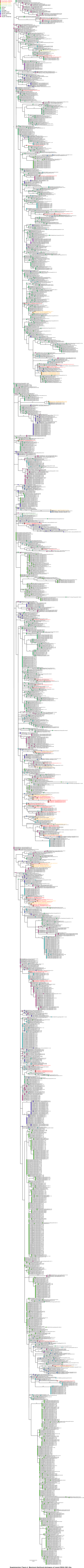


**Supplementary Figure 4. Maximum-likelihood phylogeny of recent RSVA ON1-like viruses using partial G protein sequences (n=2158)**. Branch values indicate the bootstrap replicate values with support values less 70% hidden. The boxes at each taxa reflect the region of sampling as per the key provided. The taxa labels show the GenBank accession number, Region and Country of origin, and the sampling date in YYYY-MM-DD format. The samples from this study have been colored red and orange for NSW and WA, respectively. The scale bar represents the number of substitutions per site.
