## Supplemental Table 1 for "The spatial-temporal dynamics of respiratory syncytial virus infections across the east-west coasts of Australia during 2016-17"

**Supplementary Table 1** - RSV-A clusters in NSW and WA

| Cluster | Lab | State | Lab_accession | GenBank_accession | GenBank_strain | Collection_date | Trait_age_years | Trait_location | Trait_admission | Sex | Comment |
|---|---|---|---|---|---|---|---|---|---|---|---|
| A.i | PathWest | WA | 8423521R | MW160774 | A/WA/PW8423521R/2017 | 10-May-2017 | 0.76 | SMAHS - Rockingham-Kwinana | Hospital inpatient | Female | |
| A.i | PathWest | WA | 9136191P | MW160785 | A/WA/PW9136191P/2017 | 26-Apr-2017 | 0.91 | SMAHS - Rockingham-Kwinana | Hospital inpatient | Male | |
| A.ii | PathWest | WA | 3128479T | MW160768 | A/WA/PW3128479T/2017 | 11-Apr-2017 | 0.20 | WACHS - Gascoyne | GP non-surveillance | Male | |
| A.ii | PathWest | WA | 6360608C | MW160771 | A/WA/PW6360608C/2017 | 23-Mar-2017 | 0.62 | WACHS - Gascoyne | Emergency | Female | |
| A.ii | PathWest | WA | 3128124K | MW160778 | A/WA/PW3128124K/2017 | 22-Mar-2017 | 0.62 | WACHS - Gascoyne | GP non-surveillance | Female | |
| A.ii | PathWest | WA | 7245966B | MW160783 | A/WA/PW7245966B/2017 | 17-Apr-2017 | 0.38 | WACHS - Midwest | Hospital inpatient | Male | |
| A.iii | PathWest | WA | 7225866R | MW160764 | A/WA/PW7225866R/2017 | 03-Mar-2017 | 0.98 | NMAHS - Stirling SEC | Emergency | Male | |
| A.iii | PathWest | WA | 4161683B | MW160779 | A/WA/PW4161683B/2017 | 18-Apr-2017 | 68.65 | SMAHS - Rockingham-Kwinana | Hospital inpatient | Female | |
| A.iii | PathWest | WA | 7243266W | MW160782 | A/WA/PW7243266W/2017 | 11-Apr-2017 | 1.74 | NMAHS - Oceanic | Hospital inpatient | Female | |
| A.iv | ICPMR | NSW | 20-16-276-0620 | MH760614 | A/NSW/WM0620A/2016 | 02-Oct-2016 | 20.92 | Dubbo Base Hospital | Emergency | Female | |
| A.iv | ICPMR | NSW | 20-16-276-0371 | MH760616 | A/NSW/WM0371A/2016 | 02-Oct-2016 | 0.66 | Dubbo Base Hospital | Emergency | Female | |
| A.v | ICPMR | NSW | 50-16-212-0778 | MH760630 | A/NSW/WM0778A/2016 | 30-Jul-2016 | 0.51 | Leeton Hospital | Emergency | Female | |
| A.v | ICPMR | NSW | 50-16-212-0879 | MH760631 | A/NSW/WM0879A/2016 | 30-Jul-2016 | 0.04 | Leeton Hospital | Emergency | Female | |
| A.vi | ICPMR | NSW | 40-16-274-1339 | MH760615 | A/NSW/WM1339A/2016 | 30-Sep-2016 | 63.26 | Young Hospital | Emergency | Female | |
| A.vi | ICPMR | NSW | 40-16-216-2841 | MH760626 | A/NSW/WM2841A/2016 | 03-Aug-2016 | 0.81 | Young Hospital | Emergency | Female | |
| A.vi | ICPMR | NSW | 40-16-257-2209 | MH760637 | A/NSW/WM2209A/2016 | 13-Sep-2016 | 0.71 | Young Hospital | Emergency | Female | |
| A.vi | ICPMR | NSW | 40-16-263-4313 | MH760639 | A/NSW/WM4313A/2016 | 19-Sep-2016 | 0.30 | Young Hospital | Emergency | Male | |
| A.vii | ICPMR | NSW | 03-16-272-3409 | MH760618 | A/NSW/WM3409A/2016 | 28-Sep-2016 | 1.35 | Mt Druitt Hospital | Emergency | Female | |
| A.vii | ICPMR | NSW | 03-16-271-3743 | MH760619 | A/NSW/WM3743A/2016 | 27-Sep-2016 | 75.80 | Mt Druitt Hospital | Emergency | Male | |
| A.viii | PathWest | WA | 6361561S | MW160761 | A/WA/PW6361561S/2017 | 31-Mar-2017 | 4.30 | NMAHS - Oceanic | Emergency | Male | |
| A.viii | PathWest | WA | 7245506U | MW160772 | A/WA/PW7245506U/2017 | 16-Apr-2017 | 1.40 | SMAHS - Bentley | Hospital Inpatient | Female | |
