## Supplemental Table 2 for "The spatial-temporal dynamics of respiratory syncytial virus infections across the east-west coasts of Australia during 2016-17"

**Supplementary Table 2** - RSV-B clusters in NSW and WA

| Clusters | Lab | State | Lab_accession | GenBank_accession | GenBank_strain | Collection_date | Trait_age_years | Trait_location | Trait_admission | Sex | Comment |
|---|---|---|---|---|---|---|---|---|---|---|---|
| B.i | ICPMR | NSW | 40-16-220-1032 | MH760705 | B/NSW/WM1032A/2016 | 7-Aug-2016 | 0.18 | Goulburn Hospital | Emergency | Female | |
| B.i | ICPMR | NSW | 40-16-213-2437 | MH760713 | B/NSW/WM2437A/2016 | 31-Jul-2016 | 1.50 | Goulburn Hospital | Emergency | Female | |
| B.i | ICPMR | NSW | 40-16-213-1927 | MH760717 | B/NSW/WM1927A/2016 | 31-Jul-2016 | 0.19 | Goulburn Hospital | Emergency | Male | |
| B.ii | ICPMR | NSW | 50-16-309-0066 | MH760677 | B/NSW/WM0066A/2016 | 4-Nov-2016 | 1.37 | Wagga Wagga Base Hospital | Emergency | Female | |
| B.ii | ICPMR | NSW | 50-16-279-2334 | MH760682 | B/NSW/WM2334B/2016 | 5-Oct-2016 | 1.75 | Wagga Wagga Base Hospital | Emergency | Female | |
| B.ii | ICPMR | NSW | 50-16-281-1075 | MH760690 | B/NSW/WM1075A/2016 | 7-Oct-2016 | 1.93 | Tumut District Hospital | Emergency | Male | |
| B.iii | ICPMR | NSW | 04-16-313-1775 | MH760674 | B/NSW/WM1775A/2016 | 8-Nov-2016 | 47.73 | Westmead Hospital | Hospital Inpatient | Male | |
| B.iii | ICPMR | NSW | 04-16-312-3750 | MH760676 | B/NSW/WM3750D/2016 | 7-Nov-2016 | 21.93 | Westmead Hospital | Hospital Outpatient | Male | |
| B.iii | ICPMR | NSW | 04-16-309-1924 | MH760678 | B/NSW/WM1924B/2016 | 4-Nov-2016 | 59.74 | Westmead Hospital | Hospital Inpatient | Male | |
| B.iii | ICPMR | NSW | 04-16-305-2436 | MH760680 | B/NSW/WM2436A/2016 | 31-Oct-2016 | 39.93 | Westmead Hospital | Hospital Inpatient | Male | |
| B.iv | PathWest | WA | 3543798D | MW160744 | B/WA/PW3543798D/2016 | 3-May-2016 | 0.08 | WACHS - Geraldton | Hospital Inpatient | Male | |
| B.iv | PathWest | WA | 3544838G | MW160789 | B/WA/PW3544838G/2016 | 9-May-2016 | 3.19 | WACHS - Geraldton | GP surveillance | Female | |
| B.ix | ICPMR | NSW | 20-16-229-0410 | MH760698 | B/NSW/WM0410A/2016 | 16-Aug-2016 | 90.58 | Dubbo Base Hospital | GP surveillance | Male | Nursing home outbreak of ILI |
| B.ix | ICPMR | NSW | 20-16-229-0430 | MH760699 | B/NSW/WM0430A/2016 | 16-Aug-2016 | 87.38 | Dubbo Base Hospital | GP surveillance | Male | |
| B.ix | ICPMR | NSW | 20-16-229-0425 | MH760700 | B/NSW/WM0425A/2016 | 16-Aug-2016 | 95.30 | Dubbo Base Hospital | GP surveillance | Male | |
| B.v | PathWest | WA | 3557347X | MW160745 | B/WA/PW3557347X/2016 | 26-Jul-2016 | 1.03 | WACHS - Geraldton | Emergency | Female | |
| B.v | PathWest | WA | 3555072N | MW160790 | B/WA/PW3555072N/2016 | 11-Jul-2016 | 1.09 | WACHS - Geraldton | GP surveillance | Female | |
| B.v | PathWest | WA | 3560571K | MW160791 | B/WA/PW3560571K/2016 | 15-Aug-2016 | 60.86 | WACHS - Geraldton | GP surveillance | Female | |
| B.v | PathWest | WA | 3559149Y | MW160798 | B/WA/PW3559149Y/2016 | 5-Aug-2016 | 0.05 | WACHS - Geraldton | GP surveillance | Male | |
| B.vi | ICPMR | NSW | 01-17-121-2921 | MW160813 | B/NSW/WM1212921/2017 | 1-May-2017 | 1.08 | Auburn Hospital | Hospital Inpatient | Female | Same patient |
| B.vi | ICPMR | NSW | 01-17-120-2538 | MW160835 | B/NSW/WM1202538/2017 | 30-Apr-2017 | 1.08 | Auburn Hospital | Hospital Inpatient | Female | |
| B.vii | ICPMR | NSW | 40-16-226-2459 | MH760703 | B/NSW/WM2459A/2016 | 13-Aug-2016 | 0.18 | Bega Hospital | Emergency | Male | |
| B.vii | ICPMR | NSW | 40-16-219-1630 | MH760707 | B/NSW/WM1630A/2016 | 6-Aug-2016 | 0.03 | Bega Hospital | Emergency | Male | |
| B.vii | ICPMR | NSW | 40-16-212-1183 | MH760715 | B/NSW/WM1183A/2016 | 30-Jul-2016 | 0.42 | Bega Hospital | Emergency | Male | |
| B.viii | ICPMR | NSW | 20-16-286-1234 | MH760686 | B/NSW/WM1234A/2016 | 12-Oct-2016 | 0.26 | Dubbo Base Hospital | Emergency | Male | |
| B.viii | ICPMR | NSW | 20-16-286-1228 | MH760694 | B/NSW/WM1228A/2016 | 12-Oct-2016 | 1.97 | Dubbo Base Hospital | Emergency | Male | |
| B.x | ICPMR | NSW | 03-17-095-1151 | MW160814 | B/NSW/WM0951151/2017 | 5-Apr-2017 | 0.27 | Mt Druitt Hospital | Hospital Inpatient | Female | Same patient |
| B.x | ICPMR | NSW | 03-17-096-1858 | MW160826 | B/NSW/WM0961858/2017 | 6-Apr-2017 | 0.28 | Mt Druitt Hospital | Hospital Inpatient | Female | |
