## Supplemental Figure 1 for "The spatial-temporal dynamics of respiratory syncytial virus infections across the east-west coasts of Australia during 2016-17"

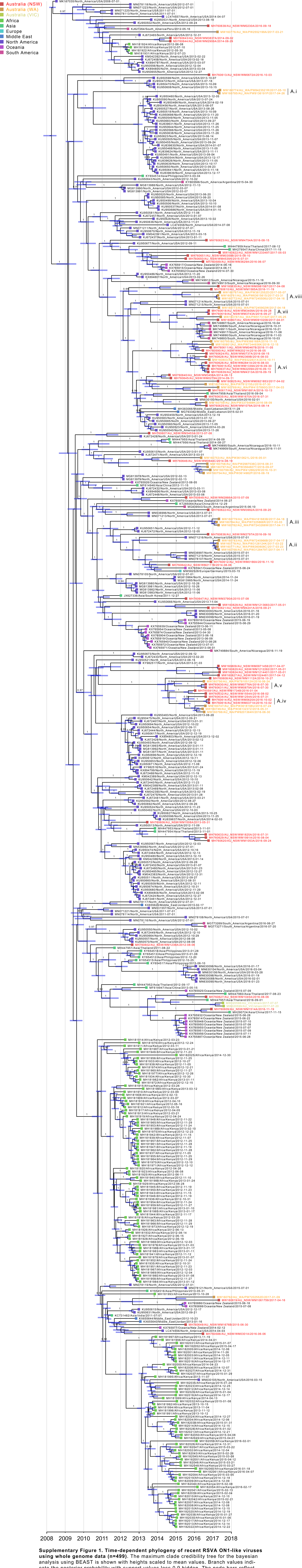


**Supplementary Figure 1. Time-dependent phylogeny of recent RSVA ON1-like viruses using whole genome data (n=499)**. The maximum clade credibility tree for the bayesian analysis using BEAST is shown with heights scaled to mean values. Branch values indicate the posterior probability with support values less 0.9 hidden. The node bars reflect the 95% HPD for height estimates. The boxes at each taxa reflect the region of sampling as per the key provided. The taxa labels show the GenBank accession number, Region and Country of origin, and the sampling date in YYYY-MM-DD format. The samples from this study have been colored red and orange for NSW and WA, respectively. The clusters identified have been labelled. The x-axis is scaled to time (year).
